# Unconscious Neural Activity Predicts Overt Attention in Visual Search

**DOI:** 10.1101/2025.02.21.639607

**Authors:** John G. Nadra, Jesse J. Bengson, George R. Mangun

## Abstract

Unconscious neural activity has been shown to precede both motor and cognitive acts. In the present study, we investigated the neural antecedents of overt attention during visual search, where subjects make voluntary saccadic eye movements to search a cluttered stimulus array for a target item. Building on studies of both overt self-generated motor actions (Lau et al., 2004, Soon et al., 2008) and self-generated cognitive actions (Bengson et al., 2014, Soon et al., 2013), we hypothesized that brain activity prior to the onset of a search array would predict the direction of the first saccade during unguided visual search. Because both spatial attention and gaze are coordinated during visual search, both cognition and motor actions are coupled during visual search. A well-established finding in fMRI studies of willed action is that neural antecedents of the intention to make a motor act (e.g., reaching) can be identified seconds before the action occurs. Studies of the volitional control of *covert* spatial attention in EEG have shown that predictive brain activity is limited to only a few hundred milliseconds before a voluntary shift of covert spatial attention. In the present study, the visual search task and stimuli were designed so that subjects could not predict the onset of the search array. Perceptual task difficulty was high, such that they could not locate the target using covert attention alone, thus requiring overt shifts of attention (saccades) to carry out the visual search. If the first saccade to the array onset in unguided visual search shares mechanisms with willed shifts of covert attention, we expected predictive EEG alpha-band activity (8-12 Hz) immediately prior to the array onset (within 1 sec) (Bengson et al., 2014; Nadra et al., 2023). Alternatively, if they follow the principles of willed motor actions, predictive neural signals should be reflected in broadband EEG activity (Libet et al., 1983) and would likely emerge earlier (Soon et al., 2008). Applying support vector machine decoding, we found that the direction of the first saccade in an unguided visual search could be predicted up to two seconds preceding the search array’s onset in the broadband but not alpha-band EEG. These findings suggest that self-directed eye movements in visual search emerge from early preparatory neural activity more akin to willed motor actions than to covert willed attention. This highlights a distinct role for unconscious neural dynamics in shaping visual search behavior.

## Introduction

Selective attention is the ability to focus one’s mental effort on behaviorally relevant objects and events while ignoring distractors. It can be directed by top-down (voluntary or goal-directed) processes (Posner, 1980) or bottom-up (reflexive or sensory) influences (Jonides, 1981), and can be done so covertly, in the absence of eye movements (e.g., Van Voorhis & Hillyard, 1977) or together with shifts of gaze as overt attention, as in visual search (Treisman & Gelade, 1980).

Studies of voluntary covert attention have benefited from trial-by-trial cue-target paradigms that temporally isolate attentional control from subsequent stimulus processing (Posner, 1978). In these experimental designs, a cue (such as an arrow) is provided by the investigator, and the observer voluntarily shifts spatial attention in line with the cue information. The cue serves as a laboratory proxy for the internal behavioral goals that drive our voluntary attention during everyday activities.

To study voluntary attention in naturalistic settings, methods have been developed to investigate self-generated attention. In such designs, rather than being cued to a location, observers are allowed to make free choices about where to direct their covert spatial attention (Bengson et al., 2014; Hopfinger et al., 2010; Taylor et al., 2008). On some trials, instead of an attention-directing cue, a free choice prompt is presented, indicating to the observer that they should freely choose where to focus covert attention on that trial. We used the term “willed attention” to describe this self-generated form of voluntary covert attention, a term introduced in our 2015 paper (Bengson et al., 2015; for a review, see Nadra & Mangun, 2023) and influenced by prior work that used the term “willed action” to refer to self-generated motor acts (Lau et al., 2004).

A key finding of these willed covert attention studies is that the pattern of brain activity in the alpha band of the EEG (8-12 Hz) in the 1 sec prior to the self-generated choice, predicts where the observer will decide to attend, a cognitive act (Bengson et al., 2015). Studies of willed motor actions have also identified predictive brain signals preceding self-paced decisions to act (e.g., reaching). In contrast to the findings in willed covert attention, in willed action, predictive brain activity has been observed in broadband EEG (Haggard & Eimer, 1999; Libet et al., 1983), rather than exclusively in the alpha-band. Also, in the studies of willed action in fMRI, it has been shown that an action could be decoded up to 8 seconds before the action (Soon et al., 2008). While research on both willed (covert) attention and willed (overt) action has revealed neural antecedents of free choices (cognitive and motor, respectively), the time course of the patterns differ significantly. The predictive brain patterns of willed attention are limited to the 1 sec prior to the free choice, while in the willed action literature, the antecedent neural activity can precede free choice motor actions by several seconds. This, along with the differing electrophysiological signals indexing each, suggest that voluntary *cognitive* “actions”, like shifting covert spatial attention, differ from voluntary *motor* actions in how they arise in the brain. But there is an interesting case where covert attention and motor actions are aligned, and that is during visual search with saccadic eye movements. Although covert spatial attention and gaze can be experimentally dissociated, in visual search tasks where attention and shifts and gaze are coordinated (de Fockert et al., 2004; Eckstein, 2011; Henderson et al., 2007; Wolfe, 2021) there is good evidence they are obligatorily linked (Belopolsky & Theeuwes, 2012). It is unknown what unconscious neural activity may precede overt (with saccades) attention during visual search.

In this report, we investigate the first shift of overt attention to the onset of a visual search array under the hypothesis that—akin to prior work on *cognitive* actions like willed covert attention (Bengson et al., 2014)—EEG alpha power patterns of brain activity in the 1 sec prior to the onset of a search array will predict the direction of the first saccade, a measure of overt attention. However, given that attention is linked to motor intention (Lau et al., 2004), and saccadic activity is a form of motor action, an alternate hypothesis is that the first saccade during visual search may arise in a fashion similar to non-saccadic willed *motor* actions, where the neural antecedents may arise much earlier (Soon et al., 2008) and be reflected electrophysiological in broadband EEG, rather than only in the alpha-band EEG (Haggard & Libet, 2001; Libet, 1985; Libet et al., 1983).

## Methods

### Participants

EEG data was recorded from 25 participants (6 males/19 females) at the University of California, Davis. The study was approved by the institutional review board. All participants had normal or corrected-to-normal vision (with contact lenses), gave informed consent, and were paid for their time. Three subjects were removed for EEG artifacts exceeding 25% of their data. Thus, the final analysis was conducted on 22 participants.

### Paradigm and Stimuli

Each trial began with a fixation point at the center of the screen. After a random 2-8 second interval, an array of triangles appeared on the screen (Figure 1). Forty potential targets spread among the field, with 24 on the lower field and 16 on the upper field. The targets were placed as though to not be detectable with covert attention when starting at the fixation location. There were also 76 distractors (19 per quadrant), which were triangles that could not be the target, but served to fill the visual field. The participant’s goal is to saccade around the visual array to find and fixate the one target (upside down triangle), then press a button to signal the end of the trial.

**Figure 1.**
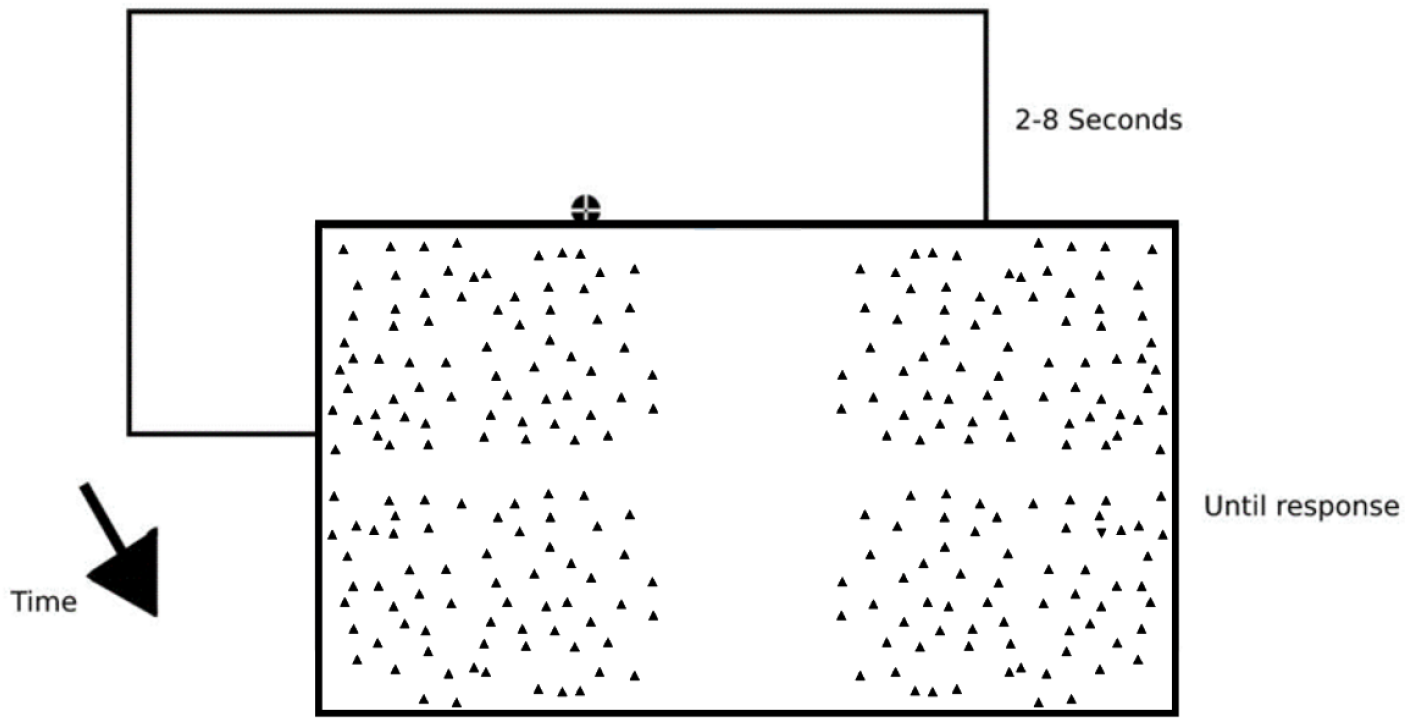
Diagrammatic representation of the visual search paradigm.

### EEG and Eye-Tracking Recording and Analysis

Eye-tracking was used to verify the subject was holding fixation before presenting the visual search array and were not blinking during the onset of the array. The eye-tracking data was collected using a Eyelink 1000 (SR Research) in remote tracking mode (target sticker affixed on forehead of participant). Nine-point calibration and validation were completed at the beginning, and drift correction was conducted between blocks periodically throughout the experiment. The EEG data was collected with a 64-channel Brain Products actiCAP active electrode system (Brain Products GmbH) and digitized using a Neuroscan SynAmps II amplifier. Eye-blink artifacts were removed via independent component analysis (ICA) as described by Vigário (1997). Residual artifacts were automatically detected, and trials with excessive artifacts were excluded using ERPLAB’s moving window peak-to-peak artifact rejection (100 μV threshold) in 100 msec windows at 50 msec intervals. Visual inspection of each trial was used to manually reject artifacts missed by automatic methods. Pre-processing utilized EEGLAB (Delorme & Makeig, 2004) and ERPLAB (Lopez-Calderon & Luck, 2014) plugins in MATLAB. Fourier analyses were conducted using the fieldtrip toolbox plugin for MATLAB (Oostenveld et al., 2011).

We employed a support vector machine (SVM) decoding pipeline similar to Bae & Luck (2018), utilizing the fitcsvm() function in MATLAB. A 3-fold cross-validated SVM was trained and tested at each time point (20 msec increments). Data were split into three equal portions, with two-thirds used for training and one-third for testing in each iteration. This process was repeated ten times, and accuracies from testing sets were averaged over iterations for each time point and trial. Nineteen relevant electrode channels (parietal and occipital) were included, and data were Fourier transformed before SVM training and testing. The binary SVM classified trials in which the first saccade landed in the left hemifield versus the right hemifield.

To assess statistical significance, we employed a nonparametric cluster-based Monte Carlo simulation, correcting for multiple comparisons. Decoding accuracy at each time point was extracted, tested using a one-sample t-test, and significant clusters (p < 0.05) were identified. Cluster t scores were compared to chance t scores from the Monte Carlo simulation to control the type 1 error rate at the cluster level. The simulation was iterated 60 times (2 bins × 3 cross-validations × 10 iterations) for each time point sampled. The data were smoothed over five time points for visualization. This process was repeated for each participant’s data. Full details about the cluster-based approach can be found in previous publications (Bae & Luck, 2018; Nadra et al., 2023; Noah et al., 2020).

## Results

### Behavioral Results

The first saccade into the visual search array landed on the left visual hemifield more than the right hemifield (55% left, 45% right – Figure 2), which aligns with studies on pseudoneglect (Durgin et al., 2008; Friedrich et al., 2018; Nicholls et al., 2017; Nuthmann & Matthias, 2014). The average latency of the first saccade into the visual search array was 316.33 ms.

**Figure 2.**
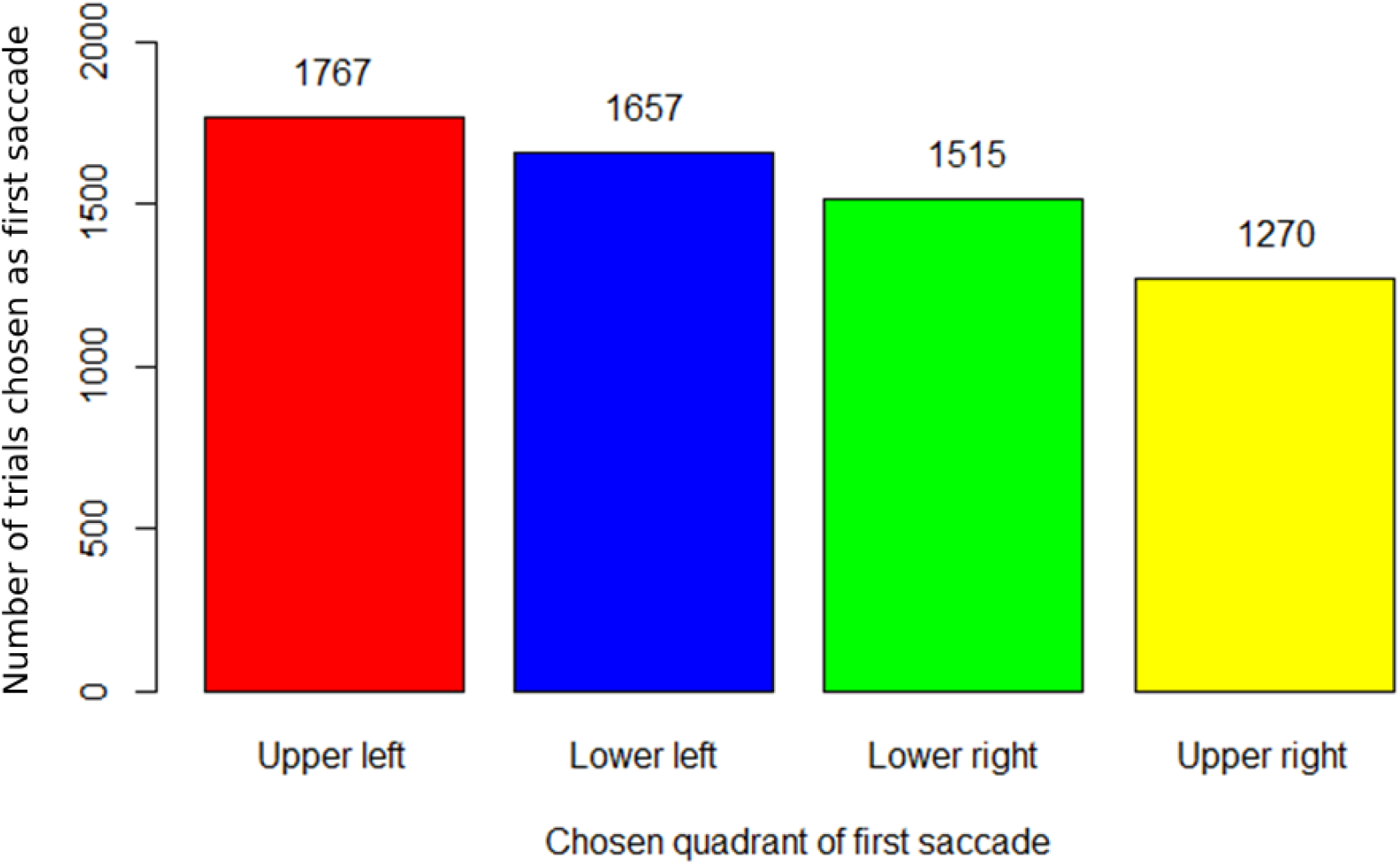
Number of trials in which first saccades landed in each quadrant of the visual field.

We computed reaction times to investigate whether the quadrant of first saccade affected search efficiency (Figure 3). There were no statistically significant differences in reaction times (RTs) as a function of the quadrant of first saccade (p > 0.05), nor were there any significant differences in RTs when collapsed across first saccades which landed in the left compared to the right visual hemifield (p > 0.05).

**Figure 3.**
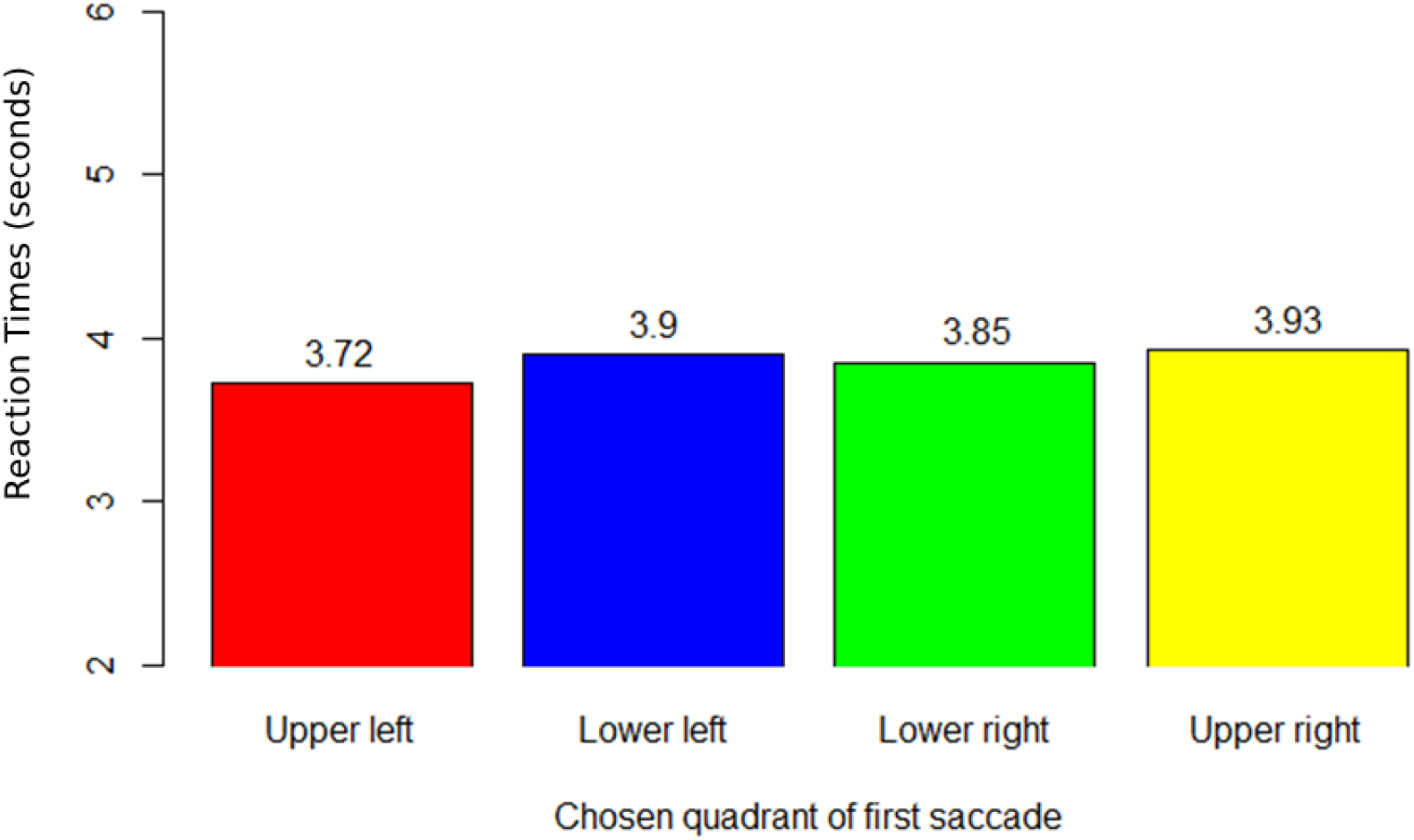
Time to detect the target (in seconds) sorted by the quadrant of the first saccade.

### Decoding Results

SVM decoding was performed on the EEG alpha power before and after the onset of the search array. Figure 4 shows the decoding accuracy from 2000 msec before the onset of the visual search array until 1000 msec after. The pattern of EEG alpha power predicted the direction of the first saccade to the search array as early as 1,200 milliseconds before array onset. This alpha pattern was time-limited, remaining significant until only about 650 milliseconds before the onset of the array.

**Figure 4.**
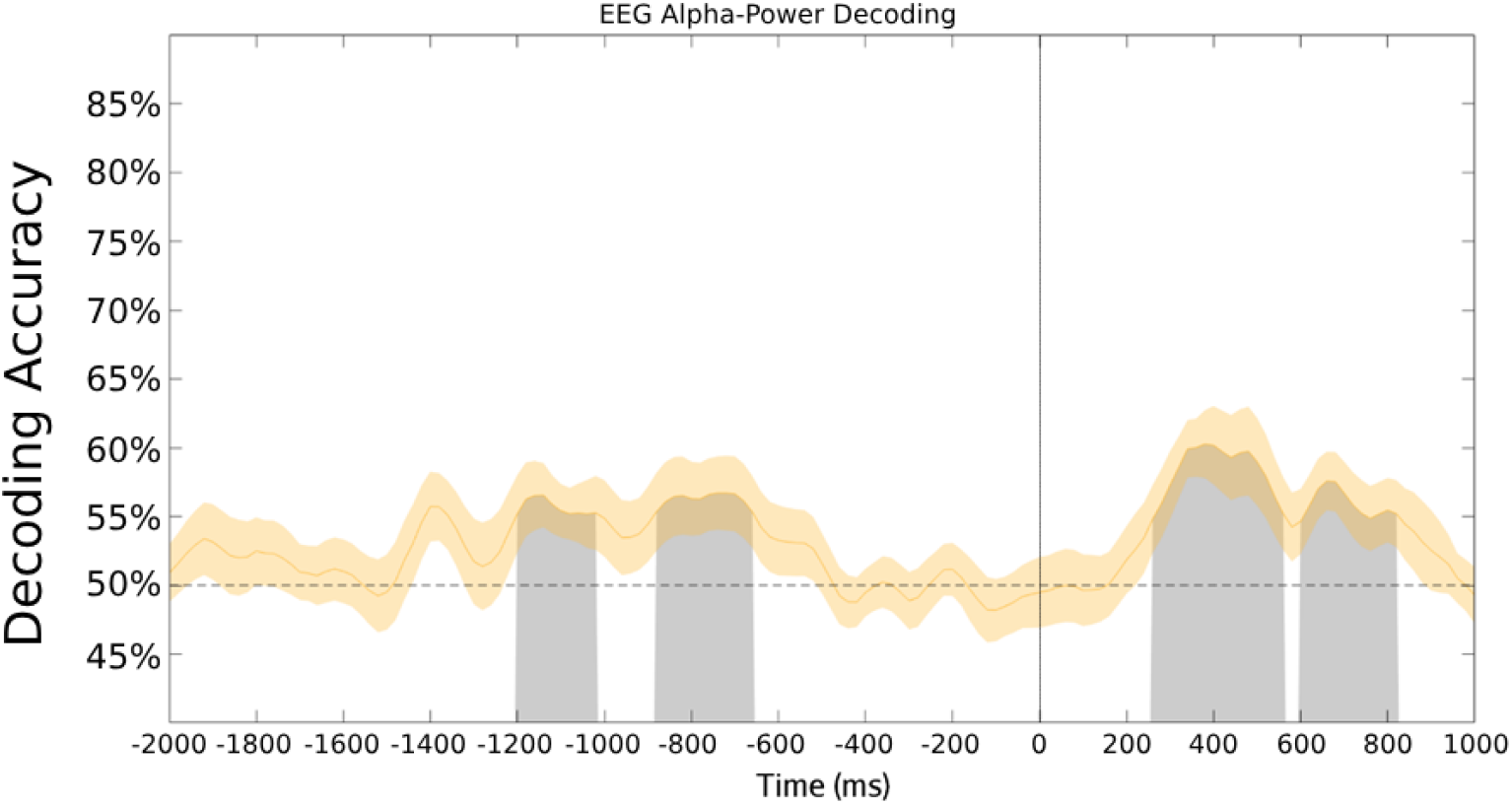
EEG Alpha Power Decoding Accuracy. Support vector machine decoding accuracy for EEG alpha power at each time point, time-locked relative to the onset of the visual search array (t=0 ms). The decoding was conducted for trials where attention was first deployed to the left compared to trials where attention was first deployed to the right. The standard error is shown as the orange shaded area surrounding the curve. Statistically significant time periods are shaded grey. This analysis was performed over 19 posterior electrodes.

The time course of predictive alpha activity in these data is quite different from that observed in prior studies of covert willed attention, where the predictive alpha signal was limited to the 1 sec immediately prior to the free choice decision about where to focus covert spatial attention (Bengson et al., 2014). Therefore, our hypothesis that overt and covert willed attention share the same underlying antecedent neural processes is not supported. To test the alternate hypothesis that overt willed attention shares mechanisms with willed motor actions, we preformed decoding analyses on broadband EEG data across the same 19 posterior electrodes (Figure 5).

**Figure 5.**
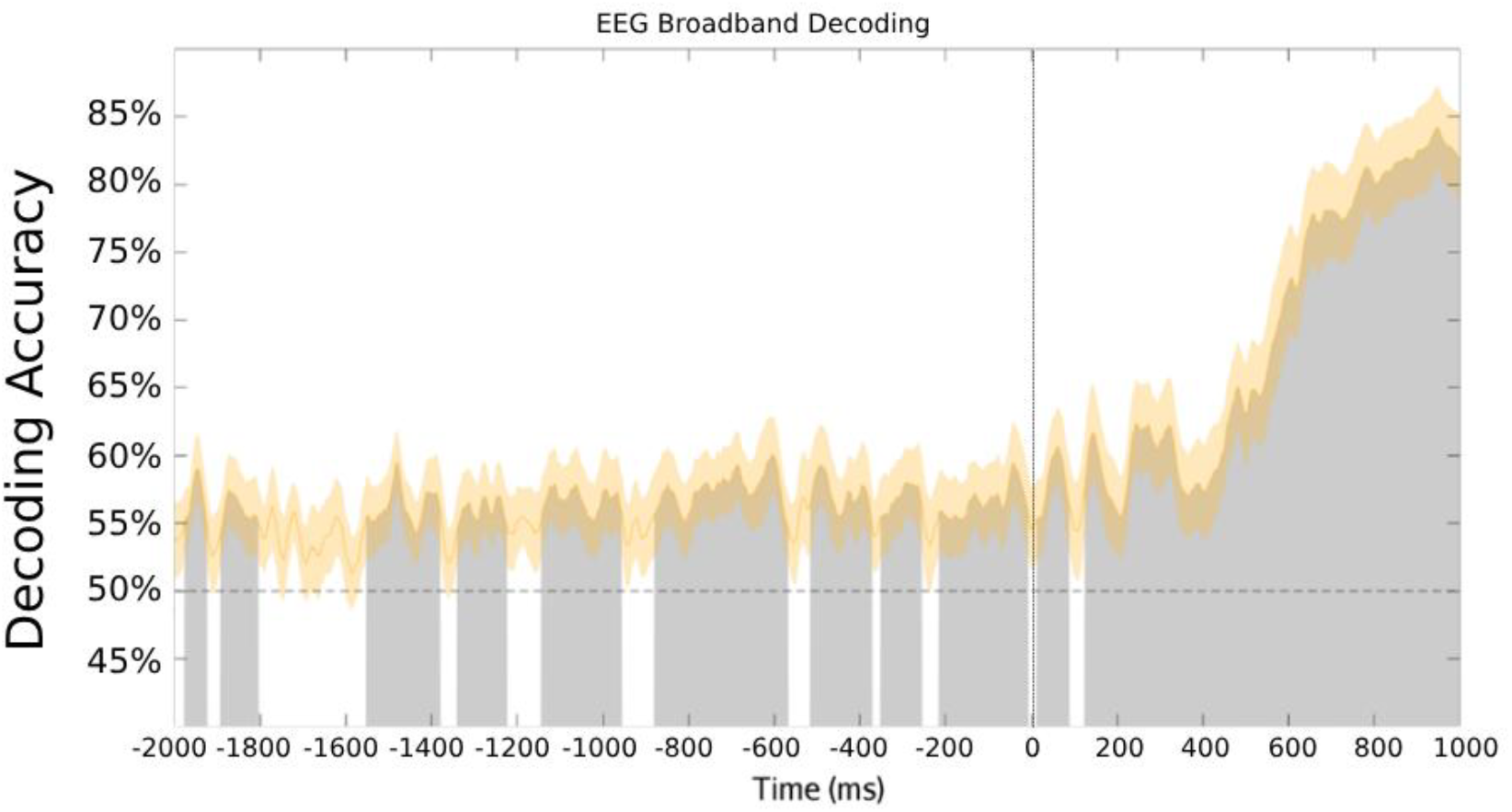
EEG Broadband Power Decoding Accuracy. As before, the decoding was conducted for trials where attention was first deployed to the left versus to the right, but now for the broadband voltage EEG data across 19 posterior channels. T=0 msec is the onset of the search array.

Figure 5 shows the decoding accuracy for the broadband EEG data. There is significant decoding throughout the whole period sampled, beginning as early as 2000 msec prior to the onset of the search array, and continuing until 1000 msec post array onset. Our statistical approach has good power and does not inflate the Type 1 error rate (Groppe et al., 2011b, 2011a) However, it is not unusual for the clusters to be separated by brief periods where the decoding dips below the uncorrected p < .05 level, as in Figure 5. This is a frequent occurrence and does not mean that the true effect is present only during the significant clusters (Sassenhagen & Draschkow, 2019). This result suggests that the neural correlates of overt willed attention are more akin to those observed in studies of willed action, in which the unconscious neural activity that predicts the upcoming action is present up to eight seconds prior to the onset of the act (Soon et al., 2008).

To test whether the significant alpha EEG decoding in the pre-array period might have been driven by the locations of the target on the previous trial (Duncan et al., 2023; Failing & Theeuwes, 2018; Talcott & Gaspelin, 2020; Theeuwes, 2019), we separated trials based on the preceding target location (left vs. right) and repeated the decoding analysis. We found no significant decoding of the current trial as a function the target location on the preceding trial (Figure 6).

**Figure 6.**
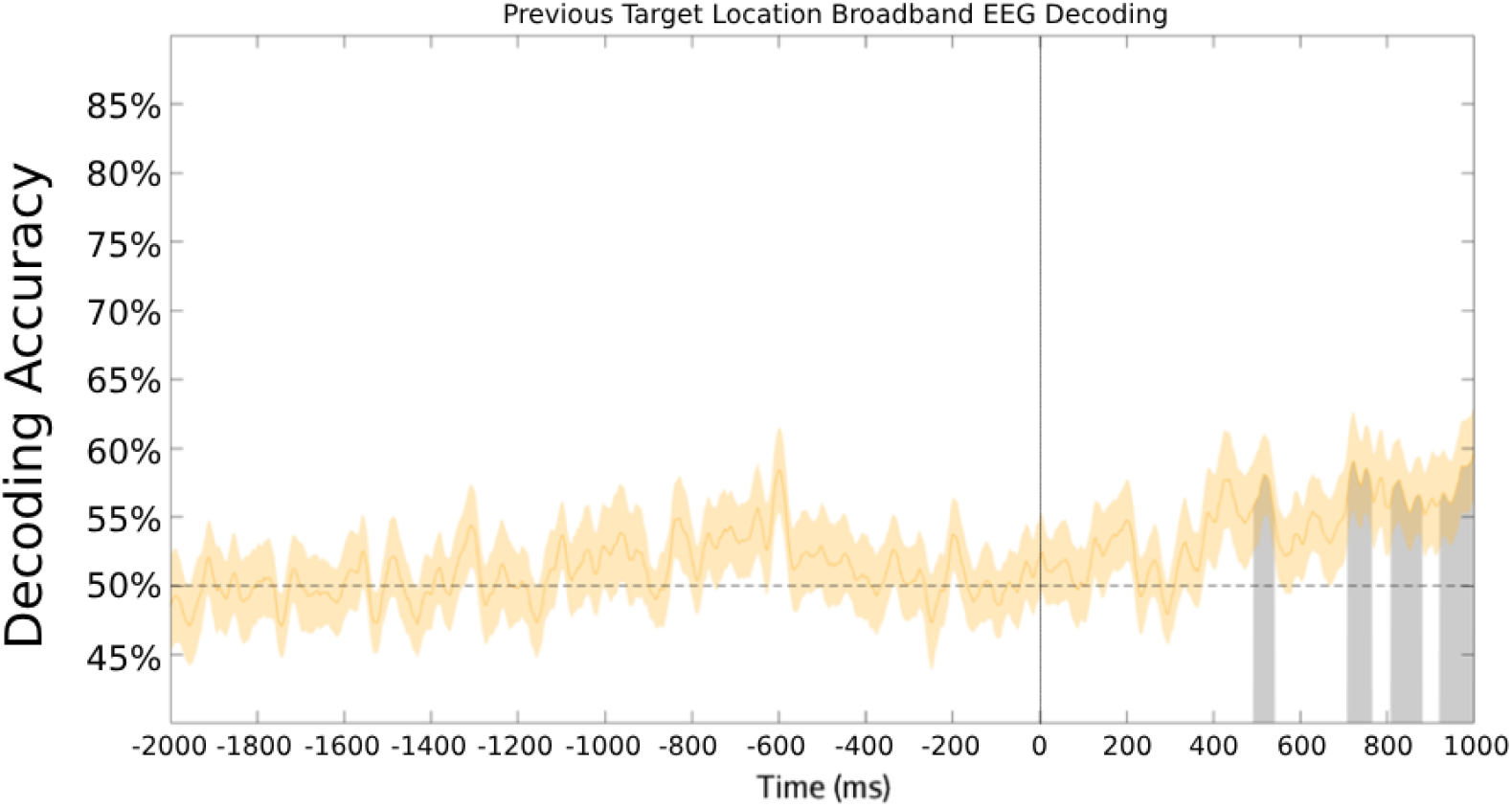
EEG Broadband Decoding as a Function of Prior Trial Target Location. Support vector machine decoding of the current trial as a function of the target location of the prior trial (left vs. right hemifield). The decoding was conducted on the broadband voltage EEG data across 19 posterior channels. T=0 msec is the onset of the search array for the current trial.

## Discussion

In this study, we investigated the unconscious neural activity that might predict the direction of self-generated shifts of overt attention (left vs. right saccade direction) at the onset of unconstrained visual search. Our analysis focused on whether the first saccade into the search array would land in the left or right hemifield, using SVM decoding to examine the period before the array’s onset. We hypothesized that the neural correlates of free choices about where to begin the search would resemble those seen in studies of covert willed spatial attention. Specifically, we expected that EEG alpha patterns during the 1 sec period before search array onset would predict the direction of the first saccade (left vs. right). An alternative hypothesis, based on the extensive literature on self-generated (willed) motor actions, predicted that brain activity would begin to predict the direction of the first saccade several seconds before the search array appeared.

Our findings support the latter prediction. Like studies of overt willed motor actions, the neural correlates of overt willed attention in visual search could be decoded from broadband EEG signals starting at least 2 seconds prior to array onset. However, contrary to our initial hypothesis, significant alpha decoding was not limited to the period immediately prior to the first self-generated saccade. These results challenge the idea that predictive antecedent brain activity in overt willed attention mirrors the literature on covert willed attention (Bengson et al., 2014). Instead, our findings align with studies of willed motor actions, where predictive brain activity can be detected well before action onset (Filevich et al., 2013; Libet et al., 1983; Soon et al., 2008, 2013). These results suggest that overt attention as reflected by saccades, may operate differently from willed shifts of covert spatial attention.

There are significant differences, though, between studies of willed motor action and willed attention, including the overt attention inherent in the current visual search study. Studies of willed motor action are typically self-paced (Hunter et al., 2003; Soon et al., 2008) without an inciting event to initiate a decision to act. Moreover, in many willed action studies, the antecedent neural activity is related only to a single action, not to a binary choice between actions. Some studies, however, have included binary options in free choice decision making across both motor (Haggard & Libet, 2001; Soon et al., 2008) and cognitive actions (Soon et al., 2013). The present study, in contrast to most studies of self-generated willed actions, used a paradigm where willed shifts of overt attention were initiated by the onset of the search array, and what was predicted by the decoding was the direction of the action, not merely its initiation. Despite these differences, the pattern of antecedent brain activity in the present study was reminiscent of the ramping up of brain activity preceding a self-generated motor action.

Our analyses also revealed that the target location on the preceding trial did not influence the direction of the first saccade in the current trial. This contrasts with other studies that have demonstrated order effects, where prior target location strongly influences the direction of first saccades (Talcott & Gaspelin, 2020). However, our average first saccade latency was 316.33 msec, which is slower than the average saccade latency in the experiments presented in Talcott & Gaspelin (2020). Factors contributing to the longer RTs may include the visually crowded search arrays we used, the fact that the array remained on the screen until participants responded, and the large, variable intertrial interval (random 2-8 sec) before the onset of our arrays. These factors could have resulted in a decay of the prior target location trace or induced strategic differences in the way participants approached the task.

Prior willed action studies in EEG have not found antecedent brain activity as early as two seconds before an action (as reported here), however, studies in EEG have shown precursor brain activity closer to the onset of the action, within 500 milliseconds (Haggard & Eimer, 1999; Libet, 1985; Libet et al., 1983). One possible reason for the present novel results is the use of machine learning methods, which are more sensitive than those used in prior EEG studies of willed action (Bae & Luck, 2018; Carrasco et al., 2023; Daly, 2023; Nadra et al., 2023; Noah et al., 2020; Trammel et al., 2023). Additional support for this idea comes from studies on willed action using pattern classification in fMRI, which found reliable decoding up to eight seconds before a decision to act (Soon et al., 2008). The timing of our decoding results aligns more closely with willed action studies in fMRI, despite the difference in temporal resolution between the two methods.

## Conclusions

In this report, we demonstrate that the direction of the first saccade during unguided visual search can be predicted by scalp-recorded EEG up to 2 sec before the unpredictable onset of the visual search array. In the context of prior studies on cognitive actions (willed covert shifts of attention) and willed motor actions (reaching or pressing a button), it is clear that our results align more closely with the findings for willed motor actions. To further investigate the unconscious neural activity influencing cognitive and motor actions, future studies should compare willed cognitive and motor actions in one experiment, analyzing both overt and covert attention using the same experimental paradigm. Overall, our results contribute to a growing body of research showing that unconscious neural activity can shape and predict behavior, offering new insights into the underlying neural dynamics of intention and action.

## Acknowledgements

This work was supported by National Institute of Mental Health grant MH117991 and National Science Foundation grant BCS 2318886. We thank Dua Haryanawalla, Lynn Fadel and Aarushi Kumar for help with data collection, and our colleagues in the Center for Mind and Brain for helpful discussions.

